# Epigenetic signatures of infection within and across generations in the endangered Loggerhead sea turtle

**DOI:** 10.64898/2026.06.25.734236

**Authors:** James O. Bazely, Eugenie C. Yen, Alice Balard, James D. Gilbert, Kirsten Fairweather, Artur Lopes, Albert Taxonera, Stephen J. Rossiter, Christophe Eizaguirre

## Abstract

Infection can substantially reduce host fitness and influence population dynamics, yet it is often difficult to detect and quantify in wild animal populations. Molecular tools offer a valuable means of identifying cryptic infection in natural systems. Using whole-genome bisulfite sequencing, we examined whether infection with the parasitic leech *Ozobranchus margoi* is associated with DNA methylation variation in loggerhead sea turtles (*Caretta caretta*), while also assessing the potential value of this variation as a biomarker of parasite infection. In nesting females, we identified infection-associated differentially methylated CpG sites associated with genes implicated in immune signalling and cellular regulation. Offspring of infected females also showed infection-associated methylation patterns, despite not being directly exposed to the parasite themselves. Differential methylation analyses identified genes involved in immunity, neurodevelopment and metabolic activity, with limited overlap in associated genes and no overlap in differentially methylated sites between generations. Maternal and offspring genome-wide methylation levels showed a non-linear association that differed subtly with maternal infection status, indicating that infection modifies intergenerational methylation associations. Finally, methylation profiles showed strong discriminatory power for maternal infection status in both maternal and hatchling samples using machine learning models, supporting their potential as candidate biomarkers of cryptic infection. Together, these results show that parasite infection is associated with distinct, generation-specific DNA methylation signatures, and highlight the potential value of epigenetic data for monitoring cryptic infection states in conservation-relevant systems.

## Introduction

Across animal taxa, parasites represent a significant selection pressure that can alter host life-history traits (Cable et al., 2017), reduce population-wide fitness (Eizaguirre et al., 2012) and, in extreme cases, contribute to population-level extinction (De Castro & Bolker, 2005). In wild populations, however, detecting and quantifying parasite infection is often challenging, particularly when symptoms are absent, mild, transient, or unevenly distributed among individuals (Grabow et al., 2024). Consequently, there is growing interest in molecular approaches that can capture an individual’s cryptic physiological response to infection in natural systems (DeCandia et al., 2018).

Given the fitness impacts of infection, hosts must mount physiological and immunological responses that are both coordinated and context dependent (O’Meara, Collins and McKenzie, 2007; Sweeny et al., 2022; Janeway, 2001). Epigenetic mechanisms can contribute to such a response by modulating gene expression without affecting the underlying DNA sequence (Dhar et al., 2021; Liotti et al., 2022; Pigliucci et al., 2006). Among these mechanisms, DNA methylation, the addition of a methyl group to cytosine to form 5-methylcytosine, has received particular attention due to its role in gene regulation and sensitivity to environmental conditions (Moore, Le and Fan, 2013; Laine et al., 2023). DNA methylation responses have been documented across a range of taxa and ecological contexts, including temperature changes (Yen et al., 2024), pollution (Baccarelli et al., 2009), and anthropogenic disturbance (Zetzsche & Fallet, 2024). In the context of parasitic infection, variation in DNA methylation has been linked to host immune responses in reptiles (e.g. Tevs, 2023), fishes (e.g. Sagonas et al., 2020), birds (e.g. McNew et al., 2021; Lundregan et al., 2022), and mammals (e.g. Al-Quraishy et al., 2013). Together, these studies suggest that DNA methylation acts as a molecular correlate of infection-related plasticity and may provide potential biomarkers for wildlife monitoring (Balard et al., 2024).

The prenatal maternal environment can also shape offspring phenotype, a phenomenon collectively referred to as maternal effects (Burgess & Marshall, 2014; Cortese et al., 2024; MacLeod et al., 2021; Thompson et al., 2018). These effects can arise through multiple pathways, including physiological state during gestation or, in oviparous species, through the provisioning of egg components such as hormones, nutrients, or other molecular signals (Agrawal et al., 2010; Groothuis et al., 2019). Maternal effects can mediate adaptive plasticity and offspring priming (Beemelmanns & Roth, 2016; Bentz et al., 2016; Kaufmann et al., 2014), as well as provide molecular information about maternal condition or exposure. This is particularly relevant for biomarker development, because maternal condition, exposure, or infection may leave detectable molecular signatures in offspring, even when offspring themselves are not directly exposed to the selection agents (Sen et al., 2015).

The extent to which DNA methylation patterns are transmitted, re-established or independently generated across generations remains heavily studied, particularly given the widespread epigenetic reprogramming which occurs during early development in many vertebrates (Bošković & Rando, 2018; Daxinger & Whitelaw, 2012). Similarities in methylation patterns between mothers and offspring may therefore reflect inherited epigenetic marks, maternally derived molecular cues, or shared developmental environments, rather than direct epigenetic inheritance per se (Heard & Martienssen, 2014). From an applied and ecological perspective, however, the utility of DNA methylation as a biomarker does not require a fully resolved causal mechanism (Balard et al., 2024): molecular signatures can be informative if they reliably predict biologically relevant traits or states, such as maternal infection history.

Understanding infection-associated molecular variation in wild species is important for developing effective conservation strategies. The loggerhead sea turtle (Caretta caretta) population in the West African archipelago of Cabo Verde provides a relevant system in which to examine such variation. This population, while one of the largest in the world (Taxonera et al., 2022), is considered endangered and has experienced an increase in the prevalence of the parasitic leech Ozobranchus margoi in recent years (Lockley et al., 2020). Although not directly lethal to sea turtles, O. margoi is associated with altered feeding ecology, reduced body condition and lower reproductive fitness, with potential consequences for host population dynamics (Lockley et al., 2020). O. margoi has also been implicated as a vector of ChHV5, the virus associated with fibropapillomatosis (Greenblatt et al., 2004; Kanat et al., 2025), further highlighting the relevance of identifying molecular correlates of infection.

In this study, we investigated whether infection with *O. margoi* is associated with DNA methylation variation in nesting loggerhead turtles, and whether maternal infection status is reflected in the methylomes of their offspring. Using whole-genome bisulfite sequencing, we tested for infection-associated differentially methylated CpG sites in mothers and hatchlings. In addition, we assessed whether maternal and offspring methylomes showed infection-dependent associations and evaluated whether infection-associated methylation profiles could serve as candidate biomarkers of maternal infection status. By combining analyses within and across generations, we aimed to assess whether DNA methylation can reveal cryptic infection-associated molecular signatures in a conservation-relevant wild vertebrate.

## Materials and Methods

### Sample Collection

Adult female loggerhead sea turtles (*Caretta caretta*) were sampled across three nights (N = 9, N = 10, N = 10) during the 2021 nesting season on the island of Sal in the Cabo Verde archipelago (Yen et al., 2026). This sampling strategy was chosen to limit incubation condition variation to track hatchling success and physiology. All samples were collected from Algodoeiro Beach (16.61773° N, −22.92882° E), an 800m stretch of sandy coastline. During oviposition, each female was visually inspected for the presence of the parasitic leech *O. margoi* and the number of parasites was counted. *O. margoi* mostly occurs on the cloaca (Göpper et al., 2018)Immediately after oviposition, 2mL of blood was collected from the dorsal cervical sinus using a 21-gauge needle and 5 mL syringe. The entire egg clutch laid by each female was relocated to an in-situ hatchery on a nearby nesting beach (16.593903° N, −22.927495° E).

Clutches were split into two equal sized sub-clutches and buried at standardised depths (35 cm and 55cm) to minimise variation in incubation conditions, which are known to influence hatchling phenology and fitness (Miguel, Anastácio and Pereira, 2022; Yen et al., 2024). Upon emergence, 100µL blood samples were collected from four hatchlings per mother, from the dorsal cervical sinus using a 26-gauge needle and 1 mL syringe. This sampling procedure was repeated for each mother-clutch pair, resulting in a total of 29 sampled mothers and 116 sampled hatchlings. Since maternal effects should be independent of hatchling incubation, we merged sub-clutches of similar maternal origin for downstream analyses. All procedures were conducted under Autorização n.°037/DNA/2021 issued by the Direção Nacional do Ambiente, Cabo Verde.

### Sample Preparation

Genomic DNA was extracted from blood samples using DNeasy Blood and Tissue Kits (Qiagen, Hilden, Germany). Library preparation for whole-genome bisulfite sequencing (WGBS) was carried out using the DNBseq method (MGI Tech, HongKong). Paired-end read sequencing (100bp) was performed on the MGI DNBSEQ platform (MGI Tech, Hong Kong). For a detailed laboratory workflow, see Yen et al. (2024).

### Methylation Data Processing and Filtering

Raw WGBS reads were quality-filtered by trimming adapters and bases with Phred scores < Q20 using cutadapt v2.10 (Martin, 2011), yielding an average of 134,716,463 ± 3,699,824 (SD) reads per sample. Trimmed reads were aligned to the chromosome-level loggerhead turtle reference genome for the Cabo Verde population (Yen et al., 2025) with Bismark v0.22.1 (Krueger & Andrews, 2011), achieving a mean mapping efficiency of 78.9 ± 3.45% (SD). Resulting BAM files were sorted and merged using SAMtools v1.9 (Li et al., 2009). Methylation calling was performed using Bismark, with a mean bisulfite conversion efficiency of 99.3 ± 0.14% (SD). CpG sites were de-stranded using the merge_CpG.py script (Cristofari, 2023), resulting in 25,182,966 ± 644,316 (SD) CpGs per sample at a mean de-stranded coverage depth of 8.91 ± 0.84 (SD). Unless stated otherwise, all downstream analysis was performed in RStudio v4.5.1 (R Core Team, 2025).

CpG sites were merged across samples using the ‘unite’ function from methylKit v1.24.0 (Akalin et al., 2012). Two independent datasets were prepared, one for mothers and one for hatchlings. For the maternal dataset, CpG sites were retained only if present in all 29 samples, to account for the relatively low sample size of this endangered species. Because the hatchling dataset is larger (N=116), CpG sites were retained if present in at least 75% of samples in the smallest treatment group (infected/uninfected) and at least the same number in the larger treatment group. For both datasets, the minimum read coverage was set at 5x and sites in the 99.9th percentile of read coverage were removed to minimise sequencing bias. Differences in sequencing depth were accounted for using methylKit’s ‘normalizeCoverage’ function. The percMethylation function from methylKit was used to generate a matrix of percent methylation at each of the retained CpG sites.

### Functional Annotation

CpG sites were annotated using the reference genome annotation developed for this population (Yen et al., 2025) using genomation v1.40.1 (Akalin et al., 2015) and GenomicRanges v1.61.1 (Lawrence et al., 2013). Promoters were defined as 1500bp upstream and 500bp downstream of a gene’s transcription start site (TSS) (Heckwolf et al., 2020). Intergenic CpG sites within 10Kb of a TSS were assigned to the nearest gene (Heckwolf et al., 2020).

For each individual in the maternal and hatchling datasets, mean methylation was calculated per genomic context (introns, exons, promoters, and intergenic regions). A linear mixed-effects model was fitted using the lme4 package v1.1.37, with methylation level as a function of the three-way interaction between generations, infection status and genomic context as fixed effects. Maternal ID was included as a random effect to account for non-independence of hatchlings. Post-hoc comparisons of estimated marginal means were conducted with *emmeans* v.2.0.0 (Lenth & Piaskowski, 2025). Benjamini-Hochberg p value correction was applied to account for multiple testing. An adjusted p value of < 0.05 was set as the significance threshold.

### Differential Methylation Analysis

To identify differentially methylated sites (DMS) between infected and uninfected mothers (DMS_M_), the *calculateDiffMethDSS* function from MethylKit was used. This function integrates a binomial mixed model (BMM) from the DSS package (H. Feng & Wu, 2019). A differential methylation analysis was also run to identify DMS between the hatchlings of infected and uninfected mothers (DMS_H_). Here, a BMM was also used using the package PQLseq v.1.2.1 (Sun et al., 2019). PQLseq integrates a genetic relatedness matrix when calculating differential methylation between groups to account for genetic structure within the data. Within this matrix, siblings from the same clutch were assigned a relatedness score of r=0.5, and unrelated hatchlings were assigned a score of r=0 (Yen et al., 2024). A q value of < 0.01 and absolute percent difference between treatments of more than 10% were set as thresholds for identifying DMS. Principal component analysis of DMS was carried out using the *prcomp* function. To test for clustering by infection status, we performed a PERMANOVA using the *adonis2* function from the vegan package v2.7.2 (Oksanen et al., 2025).

### Functional Enrichment Analysis

For both mothers and hatchlings, Gene Ontology (GO) enrichment analyses were carried out on DMS-associated genes, hereafter referred to as differentially methylated genes (DMGs). This was performed via a conditional hypergeometric GO term enrichment analysis using the R packages GOstats v.64.0 (Falcon & Gentleman, 2007) and GSEABase v.1.60.0 (Morgan et al., 2023). Two gene ‘sub-universes’ were created, one representing genes associated with hypermethylation of DMS (hyper-DMGs) and the other representing genes associated with hypomethylation of DMS (hypo-DMGs) in infected mothers and the offspring of infected mothers, respectively. These were compared against a ‘gene universe’, defined as all CpG-associated genes with available GO terms. Terms enriched in sub-universes were identified and grouped into biological processes, molecular functions and cellular components. Significant enrichment was assessed using false discovery rate with a threshold of p<0.05.

### Mother-Hatchling Association

Using 2,042,120 CpG sites where maternal and hatchling CpG methylation data overlapped, we modelled mean hatchling methylation as a function of mean maternal methylation and infection status using a generalized additive model (GAM) implemented in mgcv v.1.9.3 (Wood, 2011). At each CpG, mean methylation levels for mothers and hatchlings were aggregated across associated infection status. Smooth functions of maternal methylation were fitted separately for infected and uninfected groups to allow for infection status-specific, non-linear relationships. To accommodate for the large dataset, the model was fitted using fast restricted maximum likelihood (fREML) with discrete smoothing. To assess the contribution of infection-specific smooth terms to model performance, a model without infection-specific smooth terms was also fitted. A likelihood ratio test was then performed using these two models to assess if increased model complexity contributed to an improved model fit. This was carried out using the anova function with the “Chisq” option. To enable principal component analysis of hatchling samples at CpG identified as differentially methylated in mothers, imputation of missing values in the hatchling dataset was required. K-nearest neighbour imputation was performed using the ‘impute.knn’ function from the impute v.1.82.0 package (Hastie, 2026)

### Machine Learning for Prediction of Maternal Infection Status

We trained elastic-net logistic regression models to predict host infection status of mothers and maternal infection status in hatchlings using methylation patterns identified at DMS. Three separate models were trained using DMS_M_ values in mothers, DMS_H_ values in hatchlings, and DMS_M_ values in hatchlings, respectively. Scripts for all models were written in Python v.3.12.10. For models trained on maternal data, the StratifiedKFold function from scikit-learn v.1.80 (Pedregosa et al., 2011) was used to generate cross validation folds containing a balance of infected and uninfected samples. For models trained using hatchling data, the StratifiedGroupKFold function from scikit-learn was used to also account for genetic grouping within the data, preventing hatchlings from the same clutch appearing across both training and testing sets within a fold. KNNImputer from scikit-learn was used to impute missing values in the hatchling data. This was performed separately within each fold to minimise data leakage. The GridSearchCV function from scikit-learn was used to tune C and l1_ratio hyperparameters. Optimal values were taken forward for model validation. To assess model performance, cross-validated predictions of infection status were generated on out-of-fold samples and scores representing the probability of infection were assigned. Group-aware folds were generated for validation of models using hatchling data.

## Results

Of the 29 mothers, 20 were infected and 9 were uninfected. Parasite load ranged from one to more than 25 leeches per female. Infection status was distributed across the three sampling nights as follows: Night 1: 6 infected, 3 uninfected; Night 2: 8 infected, 2 uninfected; Night 3: 6 infected, 4 uninfected (Table S1). None of the 116 hatchlings were exposed to and therefore infected by the parasites.

### Global Methylation

Genome-wide methylation patterns were first characterised across generations, genomics contexts and infection status to assess whole-methylome variation before testing locus-specific differences. Following quality control and filtering, we retained 6,533,412 CpG sites for the 29 nesting female samples, with a mean de-stranded coverage of 11.97× (±0.39 SD). 3,573,638 CpG sites passed filtering across the 116 hatchlings with a mean de-stranded coverage of 13.15× (±0.58 SD).

Mean methylation differed significantly between generations and genomic contexts. Mothers displayed consistently higher mean methylation compared to hatchlings across all genomic contexts (exons: Δ = 6.62%, *t*_131_ = –25.976, *p* < 0.001, intergenic region: Δ = 16.38%, *t*_131_ = –64.322, *p* < 0.001, introns: Δ = 10.10%, *t*_131_ = –39.639, *p* < 0.001, promoters: Δ = 3.16%, *t*_131_ = –12.424, *p* < 0.001, Fig. 1A). The proportion of highly methylated CpG sites was also greater in mothers, with 78.33% of all sites showing mean methylation above 70%, compared to 49.71% in hatchlings.

**Figure 1:**
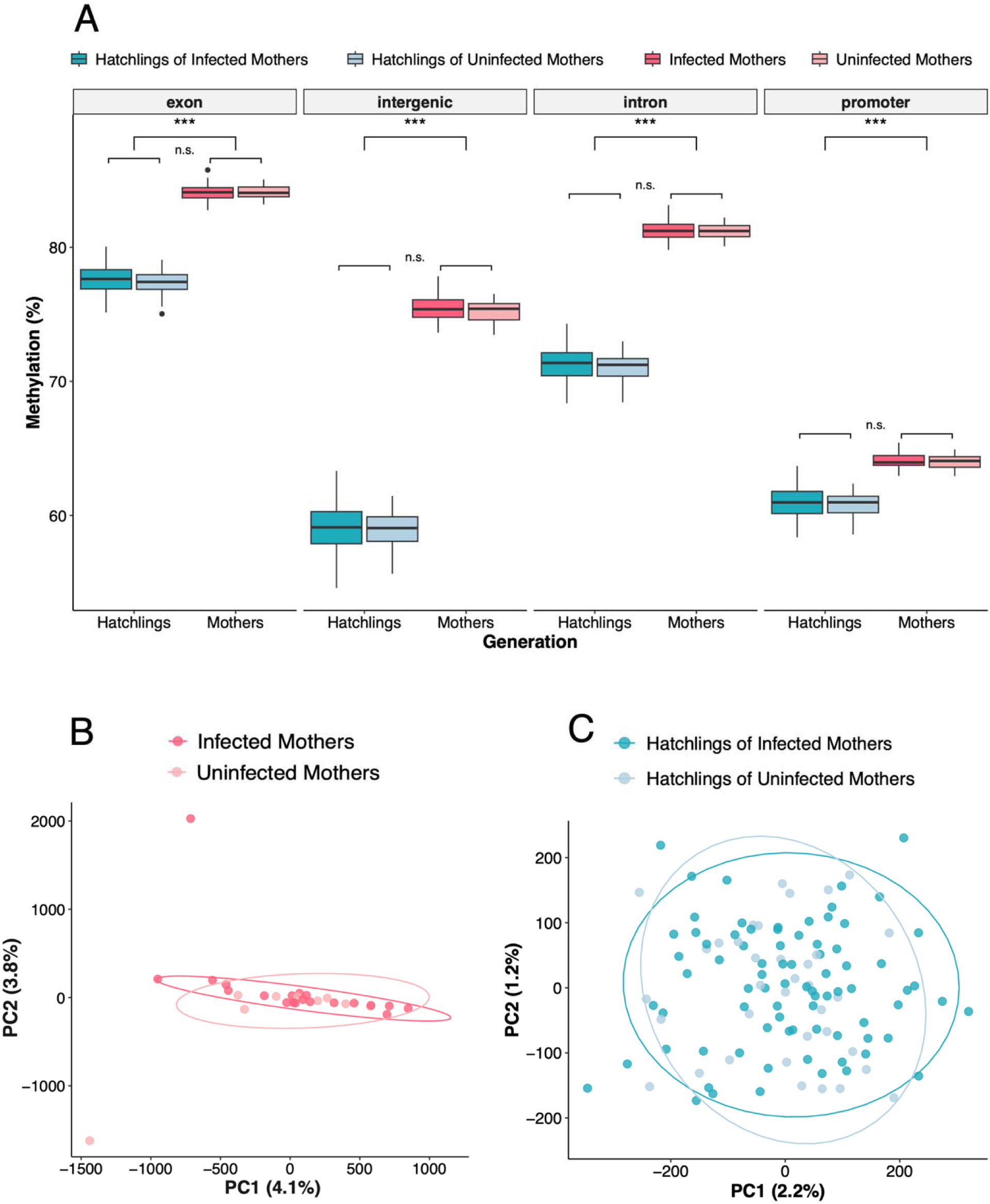
**A**) Mean methylation percentage per-generation, per feature type, split by maternal infection status. Generation (F=1210.33, p<0.001) and feature type (F=47562.01, p<0.001) Within each feature type, mothers consistently showed higher methylation than hatchlings (exon: Δ = 6.70%, intergenic: Δ = 16.55%, intron: Δ = 10.20%, promoter: Δ = 3.25%; all p < 0.001). No other comparison returned a significant difference. **B**) PCA of genome-wide hatchling methylation (n=3,573,638). **C**) PCA of genome-wide maternal methylation (n=6,533,412 CpG).

Maternal infection status was not associated with methylation levels in any genomic context or generation. When investigating patterns of global methylation, maternal methylomes did not cluster with infection status (Mothers: df = 1, F = 1.002, p = 0.302, R^2^ = 0.036, Fig. 1B). In hatchlings, on the other hand, genome-wide methylation profiles were weakly but significantly associated with maternal infection status (df = 1, F = 1.043, p = 0.041, R^2^ = 0.009, Fig. 1C).

### Differential Methylation

Differential methylation analyses were conducted separately for the mother and hatchling datasets to identify CpG sites associated with maternal infection status. In mothers, 45 differentially methylated sites (DMS_M_) were identified, with a mean absolute methylation difference of 39.80 ±7.34% (SD, Fig. 2A). Of these, 23 sites were hypermethylated and 22 were hypomethylated in infected mothers relative to uninfected mothers. 25 DMS_M_ were associated with 24 annotated genes (DMG_M_). The majority of DMS_M_ were located within intronic region of genes (n = 20), followed by exonic regions (n=2) and intergenic positions <10 kb from the nearest TSS (n = 2). A single DMS_M_ site was located in a promoter region, that of the KIAA1586 gene, which encodes for an E3 SUMO-protein ligase. SUMO E3 ligases have been implicated in immune response across a range of taxa (Qian et al., 2026; Verma et al., 2018). One DMG_M_, Rptor, was associated with two DMS_M_. The Rptor gene encodes for a subunit of the mTORC1 complex (Passtoors et al., 2013). mTOR signalling plays a key role in both innate and adaptive immune response (Chi, 2012), including response to infection in teleost fish (J. Cao et al., 2023). All other DMG_M_ were each associated with one DMS_M_ (Table S2). Principal component analysis using methylation levels at DMS_M_ showed strong clustering of mothers by infection status (PERMANOVA, df = 1, F = 14.911, p = 0.001, R^2^ = 0.356), with 36.6% of methylation variation explained by PC1 (Fig. 2C).

**Figure 2:**
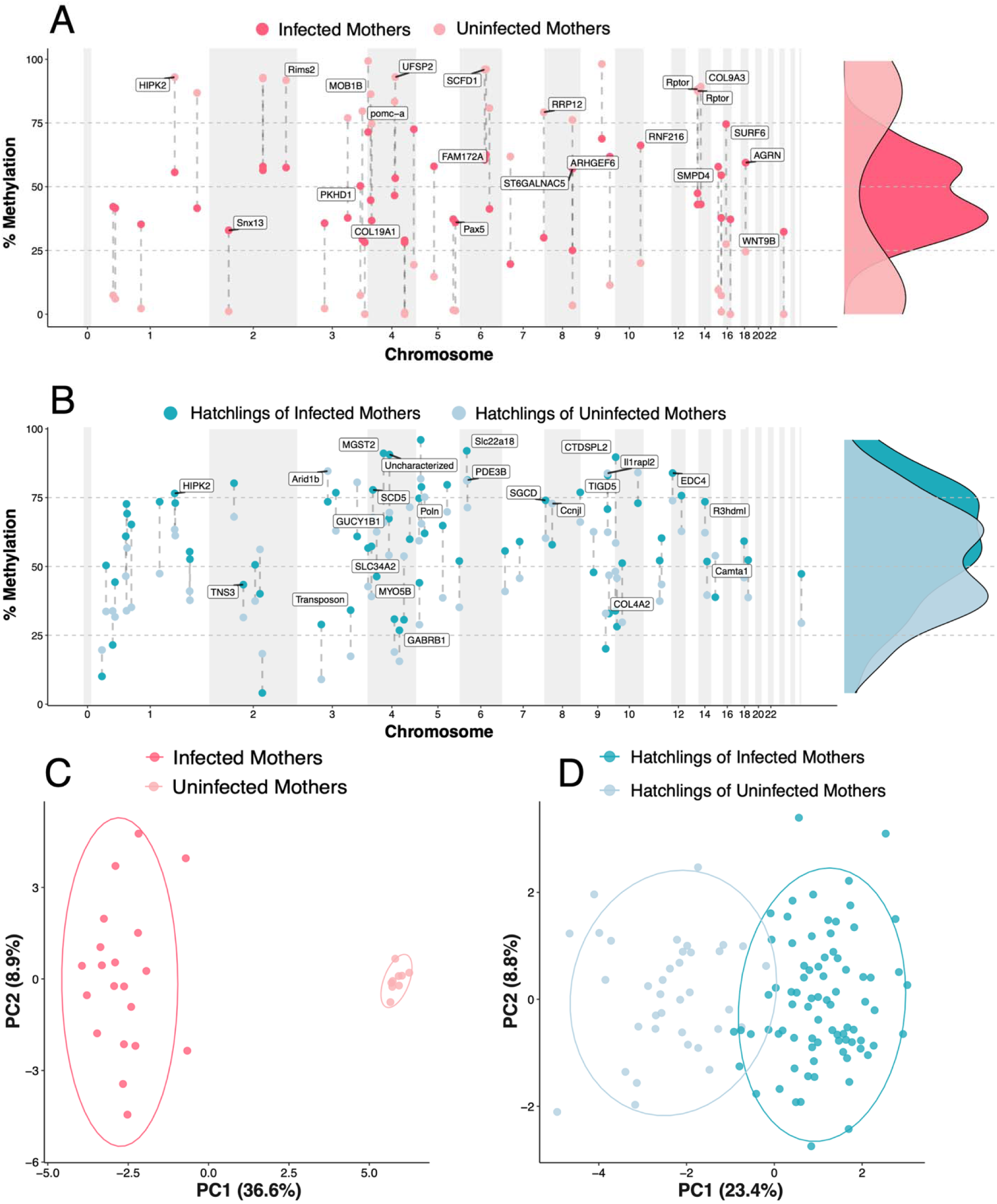
Genomic positions of differentially methylated sites (DMS) identified in **A**) mothers (n=45) and **B**) hatchlings (n=69). For each DMS, two values are shown: darker points represent mean methylation in infected individuals (or offspring of infected mothers), and lighter points represent mean methylation in uninfected individuals (or offspring of uninfected mothers). DMS associated with annotated genes are labelled with the corresponding gene name. Density plots represent infection status-specific distribution of DMS methylation levels. **C**) PC1 and PC2 co-ordinates generated by PCA of maternal methylation percentage at DMS_M_ (n=45 CpG). **D**) PC1 and PC2 co-ordinates generated by PCA using hatchling methylation percentage at subset of DMS_H_ present across all samples (n=18 CpG). For a PCA with imputed data across all DMS positions, see Fig. S2.

In hatchlings, we identified 69 differentially methylated sites (DMS_H_) between offspring of infected and uninfected mothers, with a mean absolute methylation difference of 15.27 ±5.24% (SD, Fig. 2B). Of these, 50 DMS_H_ were hypermethylated and 19 hypomethylated in hatchlings of infected mothers. A total of 30 DMS_H_ were associated with 29 annotated genes (DMG_H_), distributed across intergenic (n = 4), intronic (n = 24), and promoter (n = 2, Table S3). *HIPK2* was identified as a differentially methylated gene in both mothers (DMG_M_) and hatchlings (DMG_H_), although no differentially methylated CpG sites overlapped between generations. HIPK2 is a serine/threonine kinase whose main functions involve response to DNA damage and apoptosis (Feng et al., 2017).

Principal components analysis based on DMS_H_ methylation levels showed significant separation according to maternal infection status (df=1, F = 24.286, p = 0.001, R^2^ = 0.176), with PC1 explaining 23.4% of methylation variation at these sites (Fig. 2D). This shows a signature of maternal infection is detectable at hatchling DMS_H,_ despite hatchling not being infected with *O. margoi*.

### Functional Analysis

To assess the functional context of infection-associated DMS, we performed Gene Ontology (GO) enrichment analyses on DMS-associated genes in both mothers (DMG_M_) and hatchlings (DMG_H_) generations. In mothers, 28 GO terms were significantly enriched among DMG_M_-associated genes (Fig. 3A). Enriched categories were primarily related to signal transduction, cellular communication, mRNA regulation, and movement and secretion of molecules. GO terms with relevance to immune-related processes included TOR signalling and TORC1 complex were also identified. GTPase binding was the most significantly enriched term (p.adj < 0.001), with 12.7% of genes annotated with GTPase binding represented in the DMS-associated gene set.

**Figure 3:**
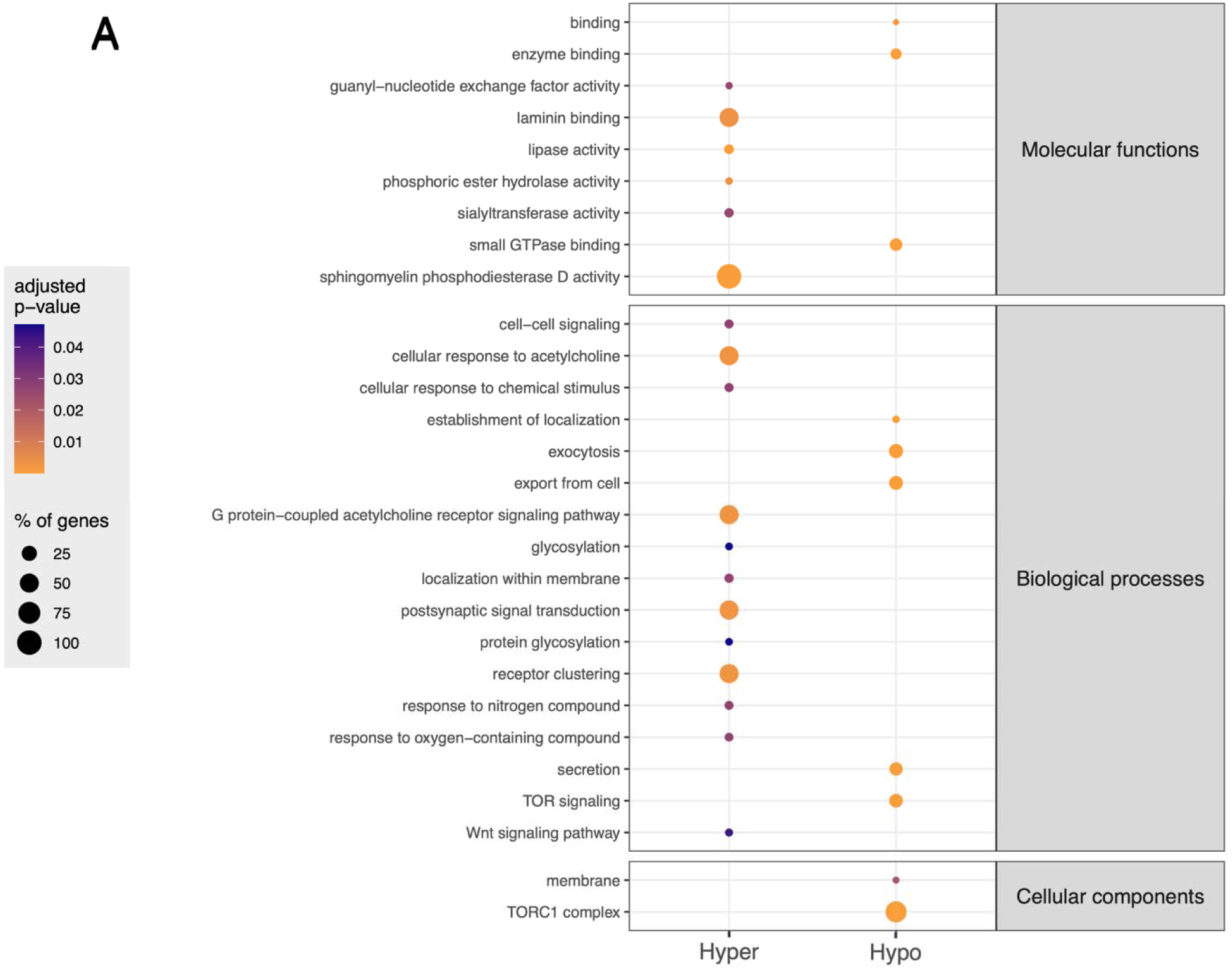
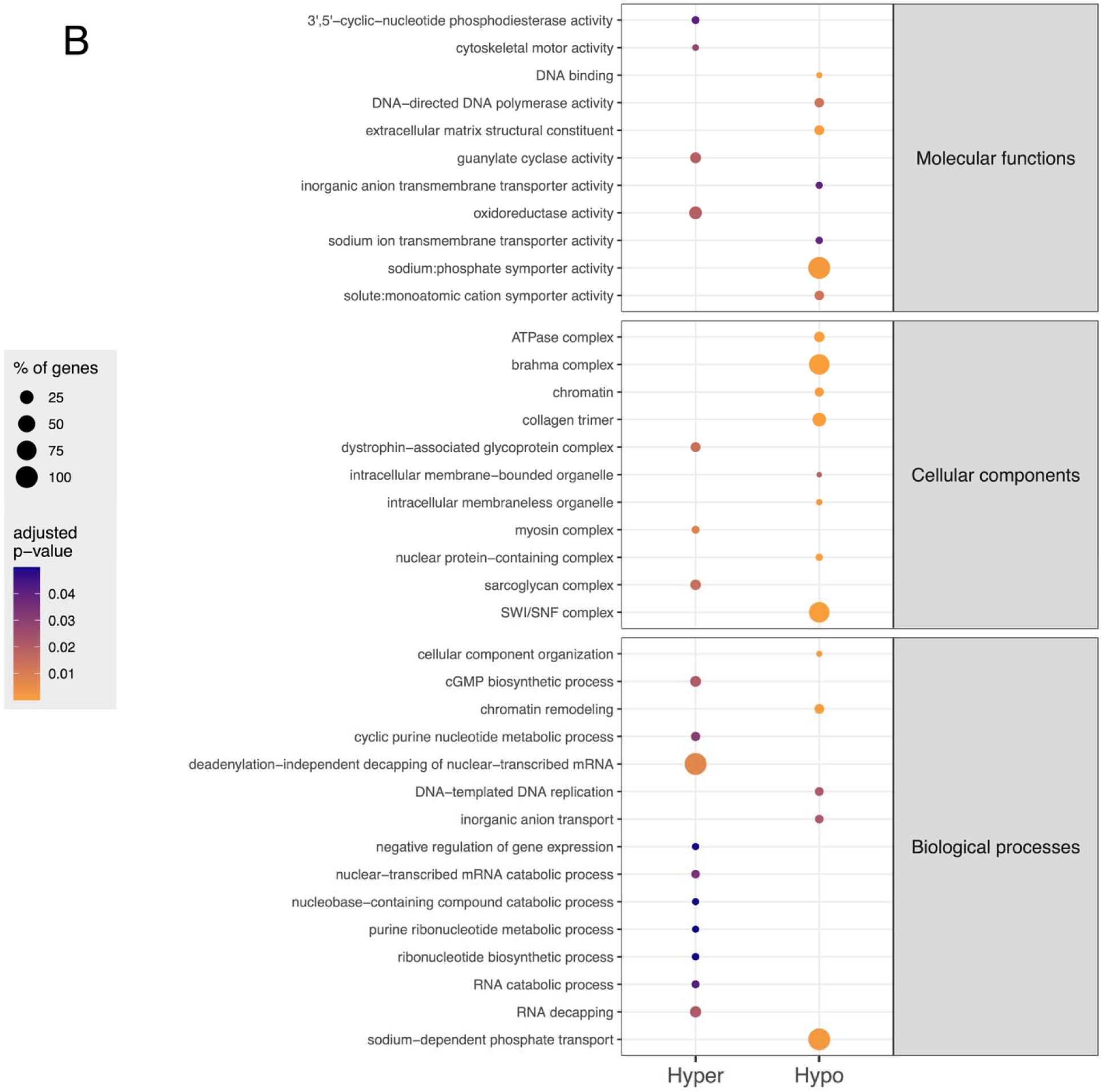
Functional enrichment of GO terms in **A**) Mothers and **B**) Hatchlings, split by association with hyper-DMG and hypo-DMG gene sets for each generation respectively. Dot size indicates the percentage of genes associated with that term that are present in the respective gene sets, and colour indicates the adjusted p-value. See supplementary tables for enriched GO terms in adults and hatchlings.

In hatchlings, 37 GO terms were significantly enriched among DMG_H_ - associated genes (Fig. 3B). Enriched categories were dominated by molecular functions related to metabolic activity, including ion channel activity, chromatin regulation, transcriptional control, and DNA replication. The most significantly enriched term was SWI/SNF complex (p < 0.001), a chromatin-remodelling complex involved in regulation of gene expression. No enriched GO terms overlapped between mother and hatchling generations.

### Intergenerational Correlation

To test whether maternal infection was associated with differences in the relationships between maternal and hatchling methylation, we fitted a generalized additive model using genome-wide methylation data from 2,042,120 CpG sites shared between generations (Fig. 4A). Across both infection groups, maternal and hatchling methylation levels showed significant non-linear associations (GAM, infected: F=648,018, p<0.001; Uninfected: F=626,034, p<0.001). Model comparison indicated that including infection-specific smooth terms significantly improved model fit (χ^2^ ≈ 1,343,016, p < 0.001), suggesting that maternal infection status is associated with a different genome-wide relationship between maternal and offspring methylomes. Although the magnitude of this effect was small (0.09% difference in mean hatchling methylation; SE = 0.012, t = 7.536, p < 0.001), it indicates a subtle but systematic infection-associated shift in intergenerational methylation association.

**Figure 4:**
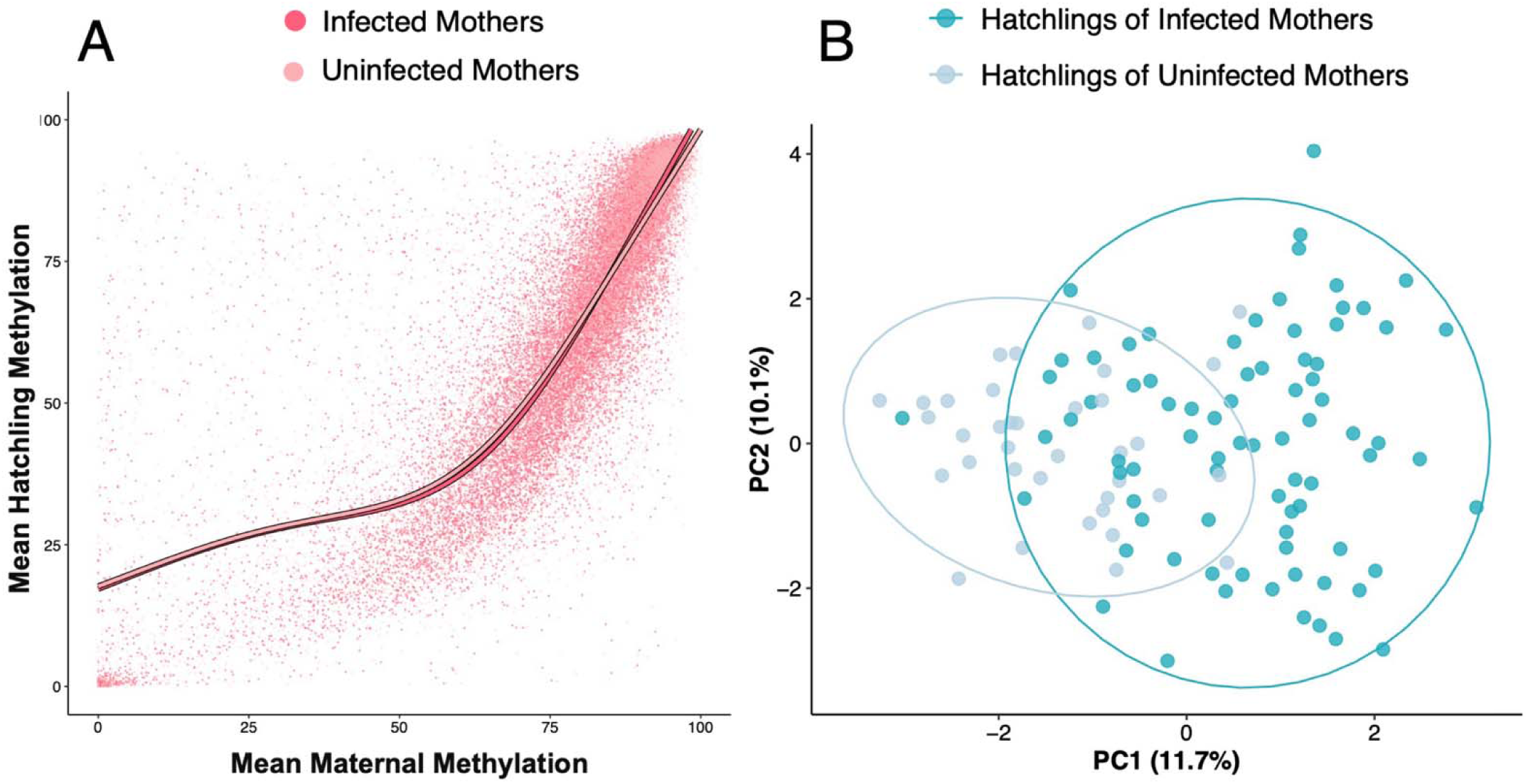
**A)** Genome-wide comparison of methylation levels at CpG present in both mothers and hatchlings by infection status (n=2,042,120 CpG). For plotting purposes, 1**%** of CpG were visualised. All data points were used when calculating and plotting smooth terms. **B)** Scatter plot of PC1 and PC2 co-ordinates generated through PCA of hatchling methylation levels at 18 maternal DMS.

Of the 45 DMS_M_ identified in mothers, 18 CpG sites were also covered in the hatchling dataset following filtering, allowing locus-specific comparison across generations. Hatchling methylation levels at these 18 DMS_M_ showed modest but significant clustering by maternal infection status (fig. 4B, F₁,₁₁₄ = 8.122, R^2^ = 0.067, p = 0.001). These results indicate that maternal infection status was associated with both a subtle genome-wide shift in mother–offspring methylation relationships and methylation differences at a subset of maternal DMS_M_ detected in hatchlings.

### DMS as Biomarkers

To evaluate whether infection-associated methylation patterns contained discriminatory information between infection status, we trained elastic net–regularised logistic regression models to predict maternal infection status using methylation at DMS_M_ in mothers, at DMS_H_ in hatchlings, and hatchling methylation at DMS_M_.

Models trained on maternal DMS_M_ and hatchling DMS_H_ respectively achieved 100% accuracy when predicting infection status in cross-validation (Fig 5A, B). In mothers, infected individuals had a mean infection likelihood score of 0.83, whereas uninfected individuals had a mean score of 0.34. In hatchlings, separation between groups was stronger, with mean infection likelihood scores of 0.92 for offspring of infected mothers and 0.21 for offspring of uninfected mothers.

**Figure 5:**
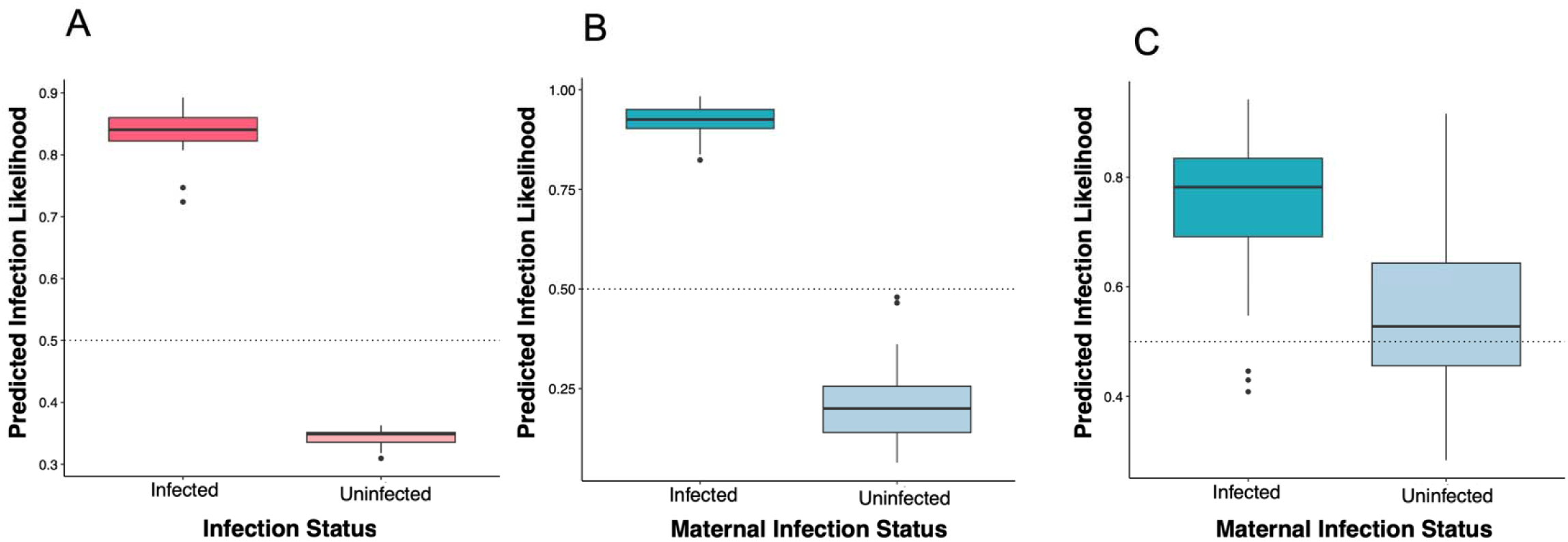
Distribution of predicted infection likelihood scores from elastic net regression models trained on **A)** Maternal methylation at DMS_M_ **B)** Hatchling methylation at DMS_H_ **C)** Hatchling methylation at DMS_M_. Y-axis represents the probability that a sample belongs to the infected (or infected mother) group based on its methylation profile, as assigned by the elastic net logistic regression models.

A third model, trained to predict maternal infection status from hatchling methylation levels at DMS_M_, achieved a balanced accuracy of 70% (Fig. 5C). This model identified hatchlings of infected mothers with a high sensitivity (96.3%) but showed lower specificity for hatchlings of uninfected mothers (44.4%). Together, these results indicate that DMS-based methylation profiles contain discriminatory information about maternal infection status in both mothers and offspring, supporting their potential as candidate biomarkers pending validation in independent samples.

## Discussion

Parasites are a strong selection pressure acting on natural populations, shaping host physiology, life-history strategies, and evolutionary trajectories. In the Cabo Verdean loggerhead sea turtle population, increasing prevalence of the parasitic leech *Ozobranchus margoi* suggests that parasite pressure may be a growing biotic challenge for a population already exposed to multiple environmental and anthropogenic stressors (Lockley et al., 2020; Yen et al., 2024). Understanding how individuals respond to infection, and whether such responses are detectable at the molecular level, is therefore relevant for both evolutionary ecology and conservation monitoring. Here, we show that *O. margoi* infection is associated with distinct DNA methylation profiles in nesting females, and that maternal infection status is also reflected in generation-specific methylation patterns in their offspring. These findings support the potential value of DNA methylation for detecting cryptic infection-associated molecular variation in conservation-relevant wild populations.

In nesting females, parasite infection was associated with differential methylation at specific CpG sites. Several DMS were located near genes (DMG_M_) with plausible roles in immunity regulation, cellular signalling and stress response. For example, the *Rptor* gene, which was associated with two maternal DMS_M_, encodes a component of the mTORC1 complex, a pathway with well-established roles in innate and adaptive immune responses (Linke et al., 2017). Other DMG_M_-associated genes, including Pax5 and WNT9B, have also been linked to immune function or cellular signalling (Calderón et al., 2021; Ljungberg et al., 2019). Gene Ontology enrichment analyses further supported an association between differential methylation and regulatory processes, with enrichment of terms including Wnt signalling, TOR signalling and the TORC1 complex. These results do not establish direct functional consequences of methylation change, but are consistent with infection-associated methylation occurring near genes and pathways relevant to host physiological responses.

Association between DNA methylation and infection have been documented across a range of host-parasite interactions. In three-spined sticklebacks, controlled parasite exposure generated genome-wide methylation differences that distinguished infected from uninfected individuals (Sagonas et al., 2020). In cattle, global methylation levels were linked to infection load with the horn fly *Haematobia irritans* (de Soutello et al., 2022), while in house sparrows, helminth infection was correlated with both genome-wide methylation differences and methylation changes at immune-related genes (Lundregan et al., 2022). Our findings extend this pattern to a naturally infected, long-lived marine reptile, showing that infection-associated methylation signatures are detectable in ecologically complex field settings, where infection may be mild, heterogeneous or difficult to quantify through external observation alone.

Probably the most central find of this study is that maternal infection status was also associated with methylation differences in hatchlings, despite the absence of direct exposure to *O. margoi*. Because *O. margoi* does not infect loggerhead sea turtles vertically, these methylation differences are unlikely to reflect direct host-parasite interactions in offspring. Instead, they suggest that maternal infection status is associated with offspring molecular phenotype, potentially through maternal effects. In oviparous species, such effects stem from egg provisioning, including hormones, nutrients, immune factors or other molecular signals, which can influence offspring developmental trajectories (Burton & Metcalfe, 2014; Marshall & Uller, 2007; Tarry-Adkins & Ozanne, 2017).

Genome-wide modelling revealed non-linear associations between maternal and hatchling methylation levels across more than two million shared CpG sites. Maternal infection status significantly modified this relationship, although the magnitude of the infection-dependent shift was small. This suggests that maternal infection is associated with a subtle but systematic change in the relationship between maternal and offspring methylomes. Importantly, however, this pattern was not accompanied by direct overlap in DMS between generations. Instead, infection-associated methylation signatures in mothers and hatchlings were largely generation-specific.

Despite the absence of shared DMS, one gene, *HIPK2*, was identified as a DMG in both mothers and hatchlings. *HIPK2* has been implicated in the production of type I interferons during antiviral immune responses (Cao et al., 2019), suggesting that related immune-associated pathways may be represented across generations. Noteworthy, the methylation differences occurred at different CpG sites and do not demonstrate direct transmission of infection-associated methylation states.

Hatchling methylation levels at maternal DMS_M_ loci also clustered significantly by maternal infection status, and methylation at these loci predicted maternal infection status with 70% accuracy using a trained elastic-net regression model. This suggests that some genomic regions carrying infection-associated methylation differences in mothers may also contain information about maternal infection status in offspring. Though tempting, evidence of an epigenetic signal at similar genomic locations across generations should not be interpreted as direct evidence of intergenerational epigenetic inheritance. Epigenetic reprogramming during embryogenesis can remodel or reset methylation marks established in parents, although the extent and timing of such reprogramming varies across taxa (Du et al., 2022; Zeng & Chen, 2019) and remains unknown in sea turtles.

A general interpretation, however, is that maternal infection may influence offspring methylation through maternal effects or developmental programming. In contexts where epigenetic reprogramming occurs during early development, maternally derived cues may contribute to the establishment of new, context-dependent methylation landscapes in offspring rather than preserving specific parental CpG states (Heard & Martienssen, 2014; Skvortsova et al., 2018, 2018). Infection can alter maternal endocrine state, metabolic allocation and immune signalling, any of which may influence egg composition and the early embryonic environment (Arshad et al., 2026; Osman et al., 2024; Widowski et al., 2022). Developmental programming of the methylome in response to embryonic environmental conditions has previously been described in the common snapping turtle (Ruhr et al., 2021). From an ecological perspective, the infection-associated intergenerational associations observed here may therefore reflect developmental plasticity, potentially shaping offspring phenotype in response to maternal condition. Equally, they could represent maladaptive maternal effects, as maternal exposure to environmental stressors can be associated with negative offspring outcomes in later life (Sheriff & Love, 2013).

Infection-associated DMS also contained sufficient signal to classify maternal infection status using elastic-net logistic regression models in both mothers and hatchlings. These results highlight candidate CpG sites with potential value as predictive biomarkers of maternal infection. Importantly, the utility of a biomarker does not depend on resolving its precise mechanistic origin; rather, its value, particularly for conservation, lies in reliably capturing biologically relevant states (Balard et al., 2024). In systems where infection may be transient, subclinical or difficult to quantify in the field, methylation-based signatures may provide complementary tools for monitoring population health.

The broader use of DNA methylation as a biomarker is already well established in the context of epigenetic clocks, where methylation patterns can reliably predict biological age across taxa (Horvath, 2013; Piferrer & Anastasiadi, 2023). These approaches also highlight the value of machine-learning models for extracting predictive information from high-dimensional epigenomic data (Libbrecht and Noble, 2015; Yousefi et al., 2022). Although sample size may be low, particularly for mothers, our elastic-net models show that infection-associated methylation profiles contain discriminatory information about maternal infection status in both generations. This therefore expands the potential scope of molecular monitoring in conservation. Where conservation genomics has traditionally provided powerful tools for assessing population structure, diversity, connectivity and adaptive potential, DNA methylation offers the possibility of predicting individual-level state or condition, including cryptic infection-associated phenotypes. In this sense, methylation-based biomarkers may contribute to precision conservation by moving from population-level genomic assessment towards individual-level inference of physiological or health status. More broadly, our findings show that maternal infection can be reflected in offspring methylomes as a predictive, generation-specific molecular signature without requiring direct inheritance of specific CpG states.

## List of Supplementary Material

**Supplementary Figure 1:** PCA of adult global methylation with outliers removed.

**Supplementary Figure 2:** PCA of hatchling methylation at DMS_H_ with missing percent methylation values imputed.

**Supplementary Table 1:** Adult metadata for sampling timepoints, infection status and number of observed *O.margoi* parasites per-individual.

**Supplementary Table 2:** DMS_M_ with associated functional annotation for gene-associated CpG.

**Supplementary Table 3:** DMS_H_ with associated functional annotation for gene-associated CpG.

**Supplementary Table 4:** Enriched GO terms for DMS_M_.

**Supplementary Table 5:** Enriched GO terms for DMS_H_.

