## Supplementary Figures for "Epigenetic signatures of infection within and across generations in the endangered Loggerhead sea turtle"

**Supplementary Material**

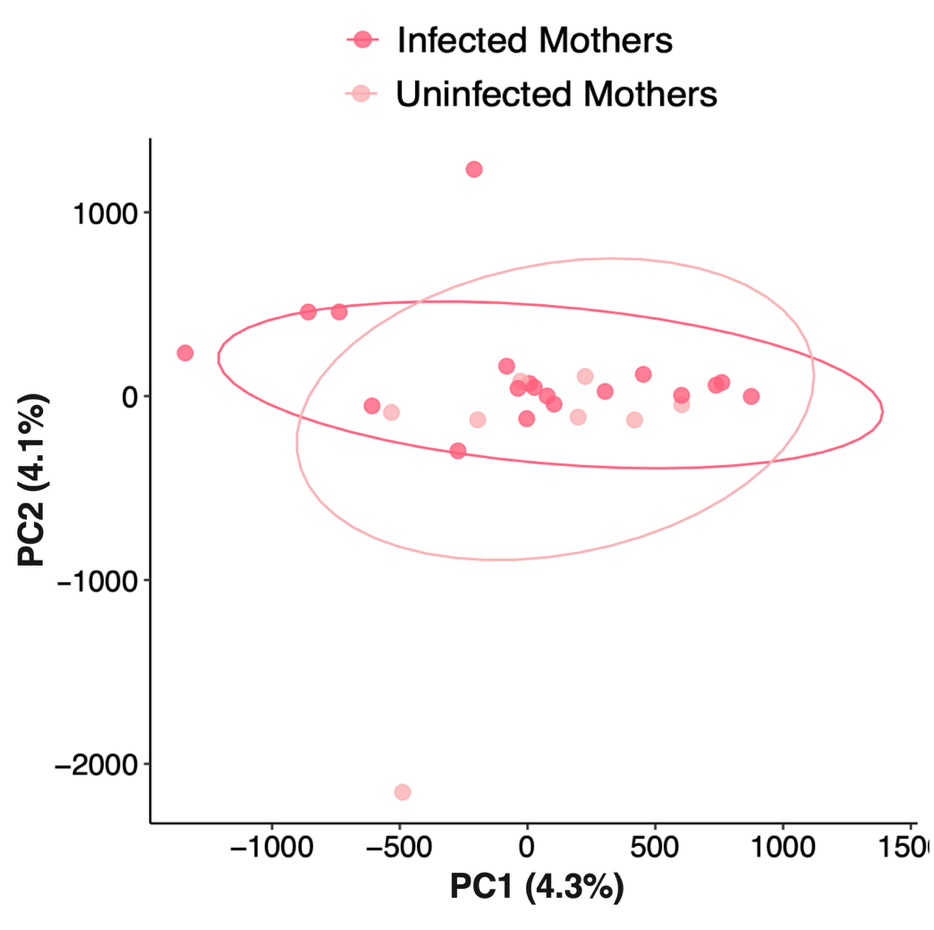

**Figure S1**) PCA of genome-wide maternal methylation following removal of two outlier samples (n=6,533,412 CpG). Samples did not significantly cluster by infection status (df = 1, F = 0.999, p = 0.452, R² = 0.038).

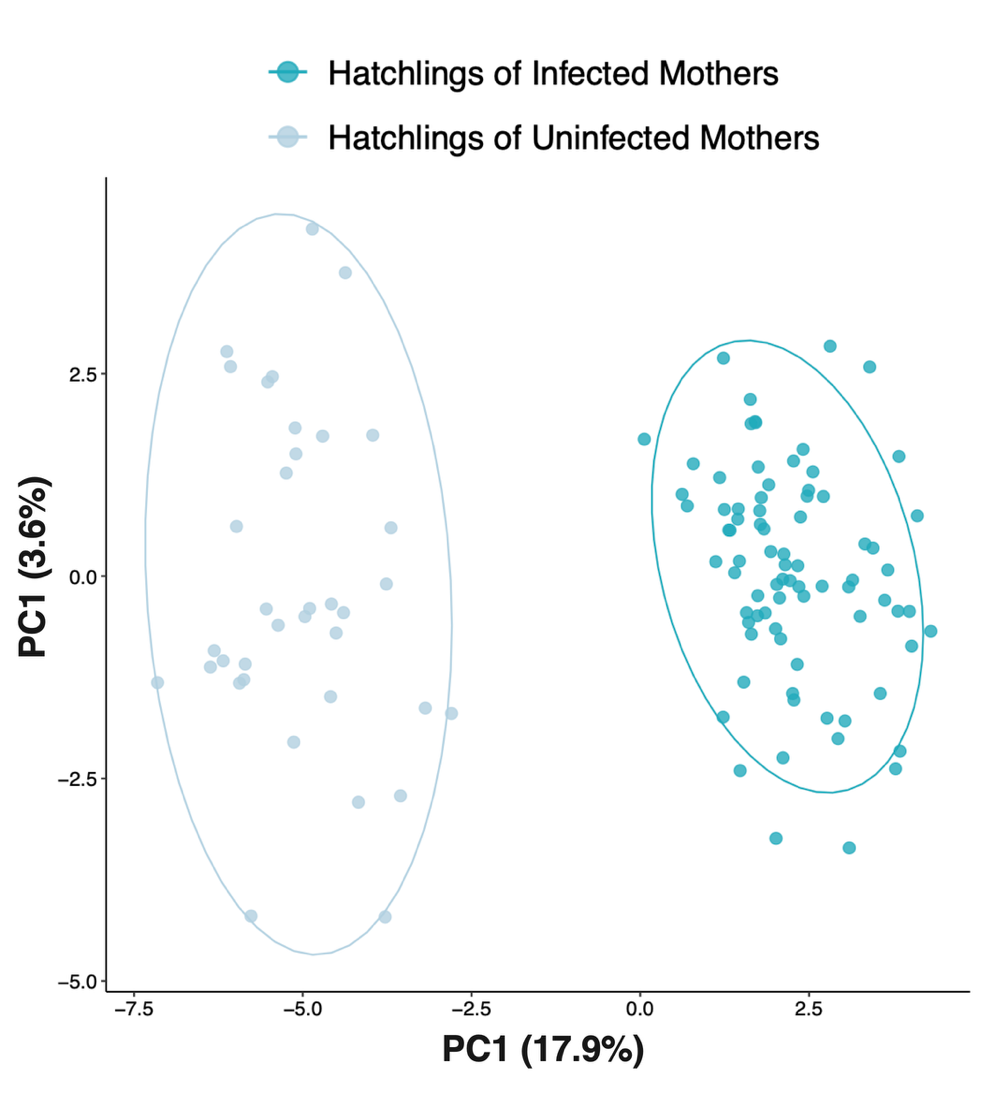

**Figure S2**) PCA of hatchling methylation at DMS_H_ with missing values imputed via k-nearest neighbour imputation (n = 69 CpG). Imputation was performed using all identified CpG in hatchlings before sub-setting for DMS_H._
